# Restriction of dietary fat, but not carbohydrate, affects brain reward regions in adults with obesity

**DOI:** 10.1101/2022.04.19.488800

**Authors:** Valerie L. Darcey, Juen Guo, Amber Courville, Isabelle Gallagher, Jason A. Avery, W. Kyle Simmons, John E. Ingeholm, Peter Herscovitch, Alex Martin, Kevin D. Hall

## Abstract

Weight loss diets often target restriction of dietary fat or carbohydrate, macronutrients that are sensed via distinct gut-brain pathways and differentially affect peripheral hormones and metabolism. However, the effects of such diet changes on human brain are unclear. We investigated whether selective isocaloric reductions in dietary fat or carbohydrate altered dopamine D2/3 receptor binding potential (D2BP) and neural activity in brain reward regions in response to visual food cues in 17 inpatient adults with obesity as compared to a eucaloric baseline diet. On the fifth day of dietary fat restriction, but not carbohydrate restriction, both D2BP and neural activity to food cues were decreased in brain reward regions. After the reduced fat diet, *ad libitum* intake shifted towards foods high in both fat and carbohydrates. These results suggest that dietary fat restriction increases tonic dopamine in brain reward regions and affects food choice in ways that may hamper diet adherence.

## MAIN

Among dietary approaches to treat obesity ^1^, popularity has waxed and waned between strategies that target restriction of dietary fat versus carbohydrate – macronutrients that elicit divergent peripheral metabolic and endocrine states ^2^. Dietary carbohydrate and fat ingestion also engages distinct gut-brain pathways affecting brain dopamine ^3-5^ which has been demonstrated in rodent models to be integral to eating behavior ^6^ and body weight regulation ^7^. While dopamine is fundamental to hedonic behaviors, the reinforcing properties of food are mediated only in part by the conscious sensory perception of pleasure *per se*. Rather, food reward is determined by signals originating predominantly from sub-conscious processes detecting nutritive cues to modulate dopamine signaling ^8^ in striatal regions involved in not only hedonic responses but also motivated behaviors, reinforcement learning, habit formation, and compulsion ^6,9^. Thus, changes in brain dopamine may affect food choice and eating behavior.

People with obesity have reduced dopamine synthetic capacity and tone ^10-12^ and availability of striatal dopamine type 2/3 receptor binding potential (D2BP) is correlated with adiposity ^13-15^. Brain dopamine has also been linked to human eating behavior ^13,16-18^ and food reward processing ^19^ independent of body weight. Whether diets restricting carbohydrates versus fat differentially affect brain dopamine and eating behavior in humans is unknown. Here, we used positron emission tomography (PET) to measure D2BP and fMRI to measure neural activity in response to visual food cues in 17 adults with obesity. We investigated whether five days of selective restriction of dietary fat or carbohydrate differentially affected brain reward regions as compared to a eucaloric baseline diet.

## RESULTS

A subset of individuals for whom metabolic results were previously reported ^2^ included 8 male and 9 female weight-stable adults with obesity, but not currently on a restrictive diet (Table 1 and Supplemental Figure 1), who had fMRI and PET neuroimaging at baseline and completed at least one of two 14-day visits to the Metabolic Research Unit at the NIH Clinical Center. As previously described ^2^, for two days prior to each inpatient admission, participants were asked to completely consume a provided standard eucaloric baseline diet (50% calories from carbohydrate, 35% calories from fat, 15% protein) that was continued for the first five days of admission. For the next six days, participants were randomized to consume a 30% calorie-restricted diet achieved either via selective reduction in fat (RF) or carbohydrate (RC), while keeping the other two macronutrients unchanged from the eucaloric baseline (Figure 1). For the final three days of each inpatient period, participants were given *ad libitum* access to vending machines stocked with a variety of supermarket foods. After a washout period of 2-4 weeks, participants were readmitted and consumed the eucaloric baseline diet for five days followed by the alternate restricted diet for six days and *ad libitum* vending machine access for three days.

**Table 1.** Characteristics of participants completing baseline and portions of neuroimaging during reduced calorie interventions.

| | Mean ± SD (Range) | N |
| --- | --- | --- |
| **Age (years)** | 34.8 ± 7.6 (23-46) | 17 |
| **Body weight (kg)** | 106.2 ± 17.2 (80.9-134.2) | 17 |
| **BMI (kg/m<sup>2</sup>)** | 36.0 ± 4.9 (29.4-44.6) | 17 |
| **% Body fat** | 39.8 ± 8.9 (22.4-51.5) | 17 |
| **Resting metabolic rate (kcal/day)** | 1826 ± 351 (1279-2323) | 17 |
| **Sex** | **Percent (%)** | **N** |
| Female | 52.9 | 9 |
| Male | 47.1 | 8 |
| **Race/Ethnicity** |  |  |
| Black | 70.6 | 12 |
| White | 11.8 | 2 |
| Hispanic | 11.8 | 2 |
| Asian | 5.9 | 1 |
| **Baseline Diet** |  |  |
| Energy intake (kcal/day) | 2685 ± 422 (2034-3399) | 17 |
| Carbohydrate intake (g/day) | 338.7 ± 52.1 (254.1-425.2) | 17 |
| Fat intake (g/day) | 104.6 ± 16.4 (79.7-129.9) | 17 |
| Protein intake (g/day) | 103.0 ± 16.3 (80.9-132.9) | 17 |
| **Reduced Carbohydrate Diet** |  |  |
| Energy intake (kcal/day) | 1993 ± 416 (1410-3133) | 17 |
| Carbohydrate intake (g/day) | 147.2 ± 29.3 (102.9-226.2) | 17 |
| Fat intake (g/day) | 110.5 ± 23.7 (78.0-175.6) | 17 |
| Protein intake (g/day) | 105.2 ± 21.1 (73.3-158.2) | 17 |
| **Reduced Fat Diet** |  |  |
| Energy intake (kcal/day) | 1909 ± 319 (1433-2385) | 15 |
| Carbohydrate intake (g/day) | 345.3 ± 57.6 (258.9-429.6) | 15 |
| Fat intake (g/day) | 15.9 ± 2.7 (12.5-21.0) | 15 |
| Protein intake (g/day) | 103.8 ± 17.1 (78.6-130.3) | 15 |

\*Reduced carbohydrate and reduced fat diets were isocaloric with subjects.

**Figure 1.**
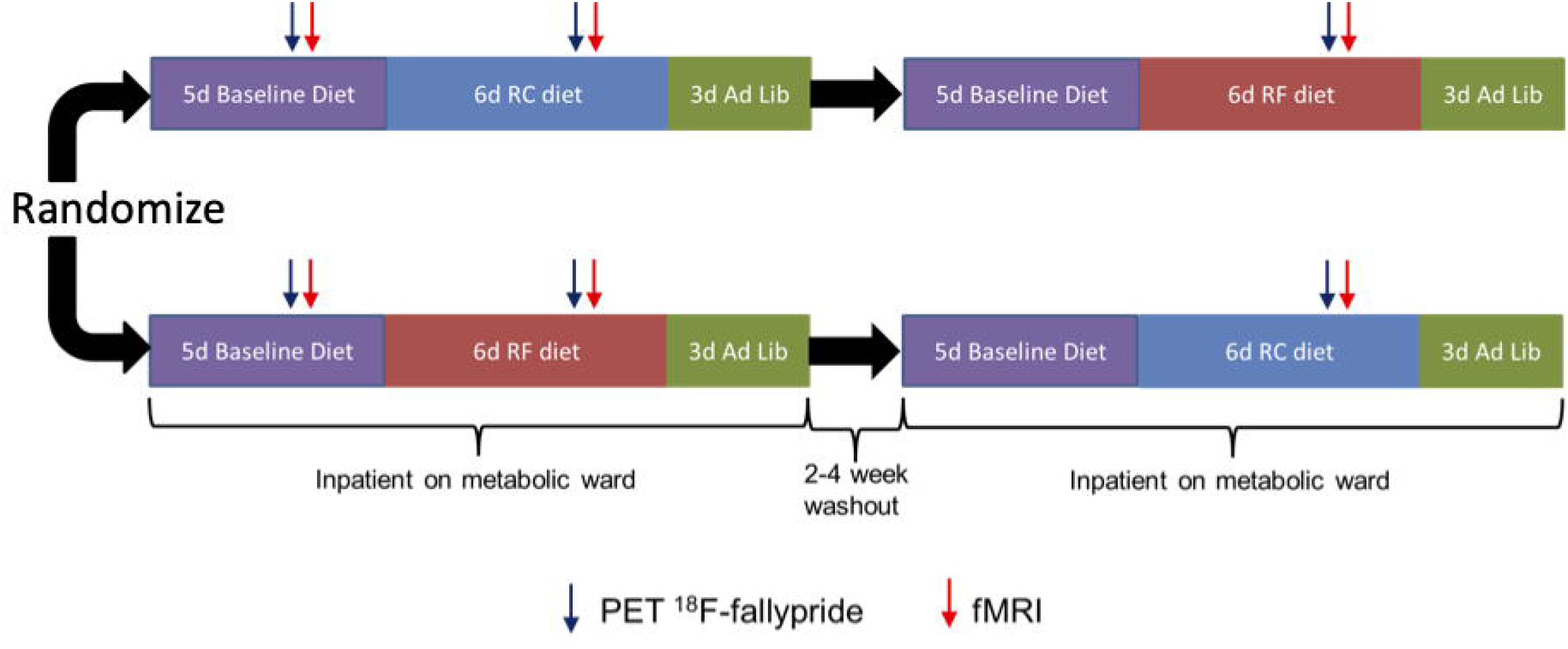
Study design. Seventeen men and women with obesity were admitted as inpatients to the Metabolic Clinical Research Unit at the NIH Clinical Center. They completed fMRI and PET scans on the third day of a five-day inpatient eucaloric baseline diet after which they were randomized to either a 30% reduced calorie diet achieved by selective restriction of dietary fat (RF diet) or carbohydrate (RC diet). Neuroimaging was repeated on the fifth day of the reduced energy diet after which, on days 12-14 of the inpatient stay, participants consumed food *ad libitum* from vending machines. After a two to four-week washout period, participants were readmitted as inpatients to complete the five-day eucaloric baseline diet and neuroimaging was repeated on the fifth day of the alternate 30% reduced calorie diet. During the final three inpatient days, participants again consumed food *ad libitum* from vending machines.

### Only the RF diet decreased activity in brain reward regions in response to food cues

Participants rated the pleasantness of a variety of food images during fMRI 4.5 hours after lunch on the fifth day (third inpatient day) of the first eucaloric baseline diet period and on the fifth day of the carbohydrate and fat restricted diets. Voxel-wise blood-oxygen-level-dependent (BOLD) responses to food images were compared to fixation within an *a priori* reward region mask encompassing orbitofrontal cortex and striatal-pallidal neurocircuit as previously reported ^20^. On average, the restricted diets did not significantly impact explicit ratings of food pleasantness in the scanner (Supplementary Figure 2). Compared to baseline, only the RF diet resulted in reduced activity in bilateral striatal clusters in caudate and putamen after correction for multiple comparisons as described in Methods (Figure 2A; Table 2). In contrast, the RC diet did not significantly change striatal responses to food images from baseline. Compared to the RC diet, the RF diet decreased activity in a dorsolateral region of the left caudate (Figure 2B; Table 2). Similar results were observed in the 15 participants with complete fMRI data for all diets (Supplementary Figure 3A-B; Supplementary Table 1), as well as the subset of 13 participants who had complete fMRI and PET data (Supplementary Figures 3C & 3D; Supplementary Table 1). Unconstrained, whole-brain analyses confirmed that the reduction in striatal activity was limited to the RF diet compared to baseline and that the RF compared to the RC diet resulted in reduced activity distributed across prefrontal clusters (Supplementary Figure 4A-B; Supplementary Table 2). Whole brain analysis of the RC diet compared to baseline revealed a significant increase in neural response to food cues in the posterior cingulate cortex (Supplementary Figure 4C; Supplementary Table 2).

**Table 2.**
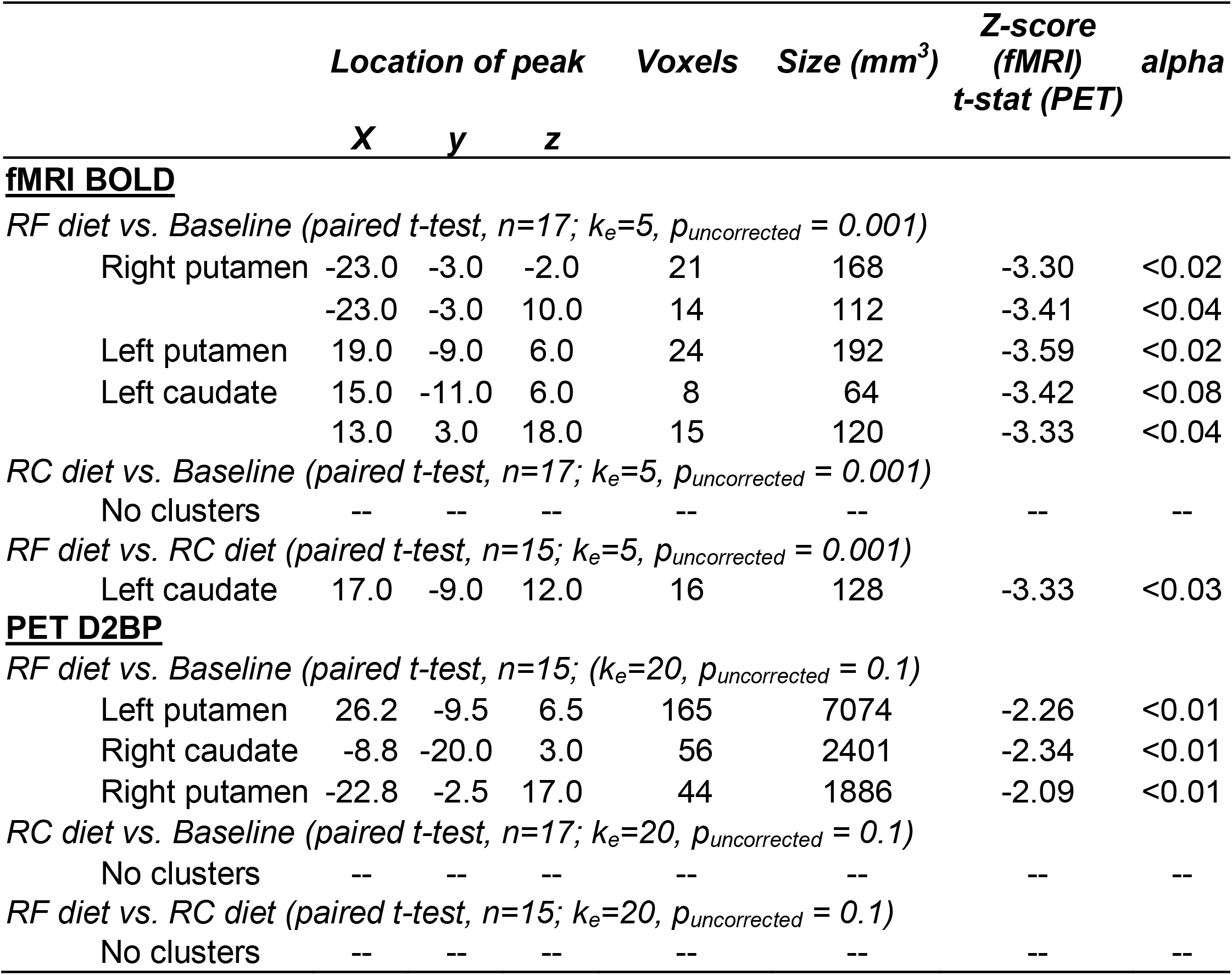
Locations of clusters displaying changes in BOLD responses within *a priori* reward region mask or D2BP to reduced fat or carbohydrate diets.

|  | Location of peak |  |  | Voxels | Size (mm<sup>3</sup>) | Z-score<br>(fMRI)<br>t-stat (PET) | alpha |
| --- | --- | --- | --- | --- | --- | --- | --- |
|  | X | y | z |  |  |  |  |
| **<u>fMRI BOLD</u>** |  |  |  |  |  |  |  |
| *RF diet vs. Baseline (paired t-test, n=17; k<sub>e</sub>=5, p<sub>uncorrected</sub> = 0.001)* |  |  |  |  |  |  |  |
| Right putamen | -23.0 | -3.0 | -2.0 | 21 | 168 | -3.30 | <0.02 |
|  | -23.0 | -3.0 | 10.0 | 14 | 112 | -3.41 | <0.04 |
| Left putamen | 19.0 | -9.0 | 6.0 | 24 | 192 | -3.59 | <0.02 |
| Left caudate | 15.0 | -11.0 | 6.0 | 8 | 64 | -3.42 | <0.08 |
|  | 13.0 | 3.0 | 18.0 | 15 | 120 | -3.33 | <0.04 |
| *RC diet vs. Baseline (paired t-test, n=17; k<sub>e</sub>=5, p<sub>uncorrected</sub> = 0.001)* |  |  |  |  |  |  |  |
| No clusters | -- | -- | -- | -- | -- | -- | -- |
| *RF diet vs. RC diet (paired t-test, n=15; k<sub>e</sub>=5, p<sub>uncorrected</sub> = 0.001)* |  |  |  |  |  |  |  |
| Left caudate | 17.0 | -9.0 | 12.0 | 16 | 128 | -3.33 | <0.03 |
| **<u>PET D2BP</u>** |  |  |  |  |  |  |  |
| *RF diet vs. Baseline (paired t-test, n=15; (k<sub>e</sub>=20, p<sub>uncorrected</sub> = 0.1)* |  |  |  |  |  |  |  |
| Left putamen | 26.2 | -9.5 | 6.5 | 165 | 7074 | -2.26 | <0.01 |
| Right caudate | -8.8 | -20.0 | 3.0 | 56 | 2401 | -2.34 | <0.01 |
| Right putamen | -22.8 | -2.5 | 17.0 | 44 | 1886 | -2.09 | <0.01 |
| *RC diet vs. Baseline (paired t-test, n=17; k<sub>e</sub>=20, p<sub>uncorrected</sub> = 0.1)* |  |  |  |  |  |  |  |
| No clusters | -- | -- | -- | -- | -- | -- | -- |
| *RF diet vs. RC diet (paired t-test, n=15; k<sub>e</sub>=20, p<sub>uncorrected</sub> = 0.1)* |  |  |  |  |  |  |  |
| No clusters | -- | -- | -- | -- | -- | -- | -- |

**Figure 2.**
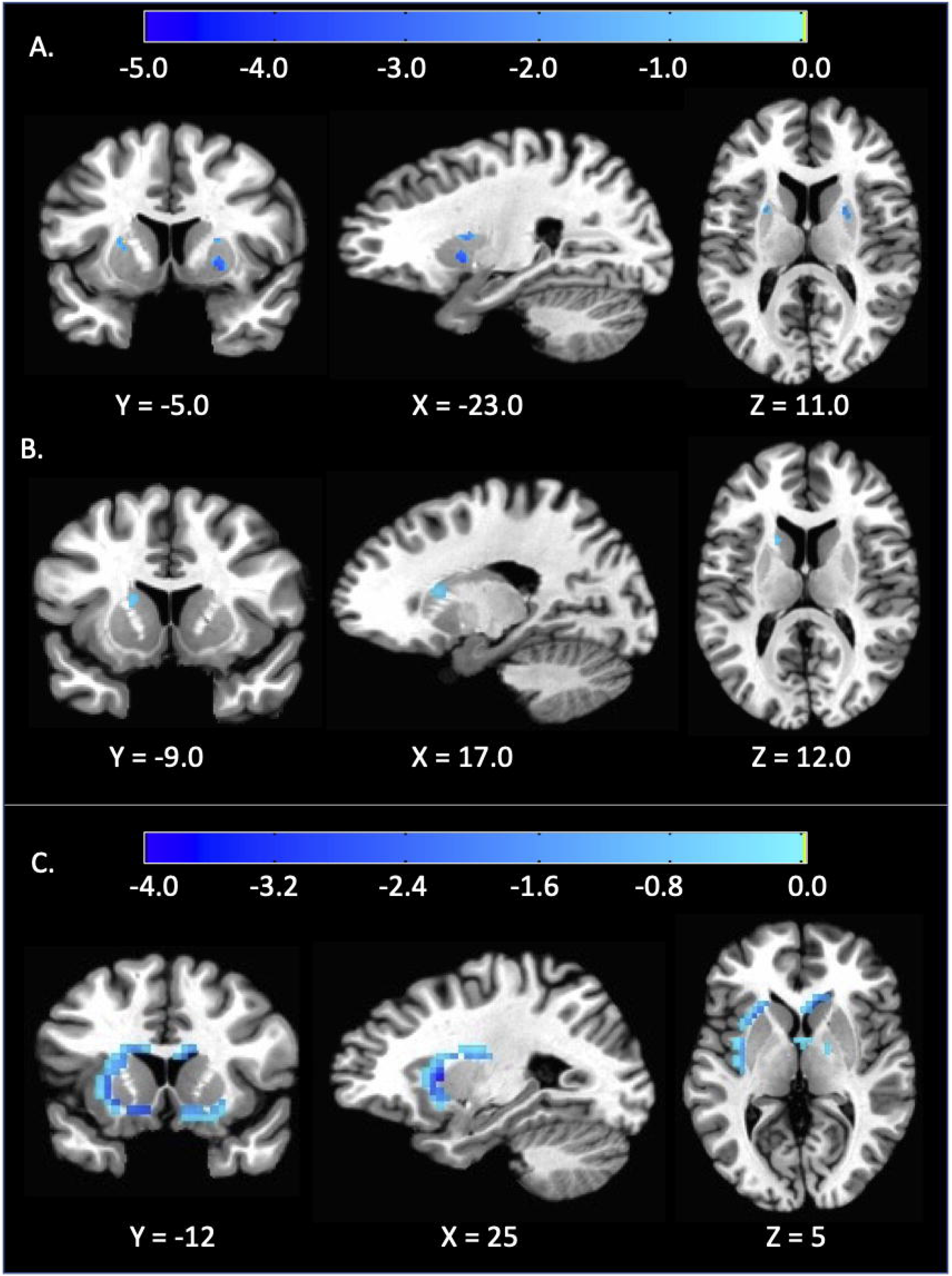
Selective reduction of dietary fat, but not carbohydrate, alters brain activity in reward regions. A) Decreased striatal response to food cues using fMRI during the RF diet compared to both the baseline diet (n=17) B) and RC diet (n=15). C) Reduced dopamine D2/3 receptor binding potential during the RF diet versus baseline using [18F]fallypride PET (n=15). Corresponding cluster details are indicated in Table 2.

### Only the RF diet led to decreased dopamine D2/D3 receptor binding potential

Participants completed PET imaging with the radiolabeled D2/3 receptor subtype antagonist [^18^F]fallypride 2 hours after breakfast on the third inpatient day as well as on the fifth day of the RC and RF diets. [^18^F]fallypride time-activity curves using the cerebellum as a reference tissue were used with kinetic modeling to measure dopamine D2/3 receptor binding potential (D2BP) as previously described ^13^. A small volume correction (D2BP > 1.5) applied to whole brain analyses was used to isolate voxel-wise D2BP analyses to the striatum.

Compared to baseline, the RF diet significantly decreased D2BP in bilateral striatal clusters spanning the left putamen and right caudate/putamen (Figure 2C; Table 2). There was no significant effect of the RC diet on D2BP as compared to baseline or the RF diet. Similar results were observed in the 15 participants with complete PET data during baseline, RF, and RC diets (Supplementary Table 1), as well as the subset of 13 participants with complete PET data who also had complete fMRI data (including three clusters surviving correction for multiple comparisons, alphas <0.05; Supplementary Figure 3E; Supplementary Table 1).

The cluster where D2BP was decreased during the RF vs. Baseline diet was localized to the white/grey matter boundary of striatal nuclei. To rule out potential localization errors due to image misalignment, individual subject alignment data was visually checked independently by two members of the study team and the mean group D2BP by diet condition was verified to map well with the template anatomical image in Talairach space (Supplementary Figure 5A). In our previously published PET processing and voxel-wise analysis pipeline ^13^, D2BP was calculated after PET data were linearly warped to Talairach space. To investigate whether cluster locations were related to the pipeline specifics, we also calculated D2BP in native space followed by non-linear warping to Talairach space. Group level BP maps for each diet condition were calculated using this alternative pipeline and indicated that peak BP signal also appropriately mapped onto striatal gray matter (Supplementary Figure 5B), but the cluster location contrasting RF and Baseline diets remained at the white/grey matter boundary (Supplementary Figure 6B).

### The RF diet resulted in greater *ad libitum* intake of foods high in both carbohydrate and fat

We explored *ad libitum* food intake for three days after the RF and RC diets. Participants selected foods from computerized vending machines stocked with calories in excess of maintenance energy requirements. Average energy intake was (mean ± SE) 25.9 ± 9.5% greater than the eucaloric baseline diet and was not significantly different following RF versus RC diets (Table 3). While overall macronutrient intake was similar after RC and RF diets, participants consumed a greater percentage of total calories from foods high in both carbohydrate and fat (HCHF) as well as high in both sugar and fat (HSHF) following the RF diet as compared to the RC diet (RF diet 28.8 ± 2.4% vs. RC diet 23.1 ± 2.4%; p=0.010) and consumed more calories from sugar sweetened beverages (SSB) (RF diet 9.8 ± 1.1% vs. RC diet 8.4 ± 1.1%; p=0.032) such that the combination of HCHF, HSHF, and SSB as a fraction of total calories consumed was greater following the RF diet (RF diet 38.6 ± 3.0% vs. RC diet 31.4 ± 3.0%; p<0.001).

**Table 3.** *Ad libitum* intake over 3 days from vending machines after RF and RC diets (mean ± SE; n=17).

|  | **After RC** | **After RF** | ***p*** |
| --- | --- | --- | --- |
| ***Ad Libitum* Energy Intake (kcal/d)** | 3225 ± 306 | 3297 ± 306 | 0.629 |
| **Protein (%kcal)** | 16.5 ± 0.9 | 16.0 ± 0.9 | 0.315 |
| **Fat (%kcal)** | 39.6 ± 1.2 | 39.3 ± 1.2 | 0.770 |
| **Carbohydrate (%kcal)** | 44.3 ± 1.1 | 45.4 ± 1.1 | 0.365 |
| **Sugar, total (%kcal)** | 20.0 ± 0.9 | 21.0 ± 0.9 | 0.231 |
| **Items selected per day (total)** | 19.3 ± 1.2 | 19.9 ± 1.2 | 0.427 |
| **Ultra-processed foods (%kcal)** | 80.7 ± 2.3 | 81.2 ± 2.3 | 0.748 |
| **Hyperpalatable foods (%kcal)** | 70.7 ± 2.8 | 71.2 ± 2.8 | 0.762 |
| **HCHF foods (%kcal)** | 9.8 ± 1.7 | 12.8 ± 1.7 | 0.072 |
| **HCLF foods (%kcal)** | 11.9 ± 1.5 | 10.2 ± 1.5 | 0.246 |
| **HPHF foods (%kcal)** | 25.5 ± 2.0 | 21.7 ± 2.0 | 0.049 |
| **HPLF foods (%kcal)** | 10.2 ± 1.6 | 11.4 ± 1.6 | 0.429 |
| **HSHF foods (%kcal)** | 13.2 ± 1.5 | 16.0 ± 1.5 | 0.099 |
| **HSLF foods (%kcal)** | 6.6 ± 1.3 | 5.5 ± 1.3 | 0.159 |
| **SSB (%kcal)** | 8.4 ± 1.1 | 9.8 ± 1.1 | 0.032 |
| **HCHF+HSHF (%kcal)** | 23.1 ± 2.4 | 28.8 ± 2.4 | 0.010 |
| **HCHF+HSHF+SSB (%kcal)** | 31.4 ± 3.0 | 38.6 ± 3.0 | <0.001 |

HCHF=high carbohydrate, high fat; HCLF=high carbohydrate, low fat; HPHF=high protein, high fat; HPLF=high protein, low fat; HSHF=high sugar, high fat; HSLF=high sugar, low fat; SSB=sugar sweetened beverages

## DISCUSSION

We previously showed that the RC diet led to widespread metabolic and endocrine changes compared to the eucaloric baseline diet, including increased fat oxidation as well as decreased energy expenditure and decreased daily insulin secretion, whereas the RF diet did not lead to substantial peripheral metabolic or endocrine changes ^2^. Therefore, we expected that the RC diet would have a greater effect on brain reward regions than the RF diet, especially given insulin’s effects on dopamine levels ^21,22^. Surprisingly, it was the RF diet, and not the RC diet, that significantly decreased both D2BP and neural activity in response to visual food cues in brain reward regions as compared to the baseline diet. The fMRI data showed decreased activity in brain reward regions in response to visual food cues during the RF diet as compared with the RC diet, but there were no significant differences in D2BP between RC and RF diets. Furthermore, *ad libitum* food intake after the RF diet was shifted towards high fat, high carbohydrate foods as compared to the RC diet. These results suggest that a calorie is *not* a calorie when it comes to macronutrient effects on brain reward regions in humans.

The most likely interpretation of our data is that the RF diet increased striatal tonic dopamine. This would explain the observed decrease in D2BP because increased endogenous dopamine would be expected to displace the [^18^F]fallypride tracer ^23,24^. Furthermore, an increase in tonic dopamine would be expected to activate high affinity D2/3 receptors thereby inhibiting neural activity ^25^ and explaining the observed decrease in brain activity to visual food cues with the RF diet. Indeed, pharmacological agonism of the D2/3 receptor has been demonstrated to decrease the fMRI signal in both rats ^26^ and non-human primates ^27^.

D2 receptors are located both post-synaptically on non-dopaminergic cells within the striatum and pre-synaptically on cell bodies, axons and axon terminals of dopaminergic projection neurons ^28^. D2 receptors found pre-synaptically function as autoreceptors to modulate dopamine signaling ^28^ and are of relevance to control of human behavior ^29^. The RF diet decreased D2BP at the white/gray matter boundary of striatal nuclei, with a peak in apparent white matter which therefore might indicate differences in endogenous dopamine acting pre-synaptically on dopamine autoreceptors. Alternatively, localization to this region might have simply resulted from the limited resolution of PET to detect differences in D2BP at the edge of the striatum ^30^.

It is unlikely that the observed reduction in D2BP during the RF diet was due to decreased D2/3 receptor density because neither dopamine depletion over 2-5 days ^24,31^ nor dopamine stimulation over five weeks ^32^ appreciably impacts receptor density. Moreover, a reduction in D2/3 receptor density would be expected to both minimize D2/3 receptor inhibitory signaling and produce a net increase in the stimulatory effect of dopamine at neurons expressing D1/5 receptors ^27^ which is inconsistent with the observed decrease in fMRI response during the RF diet. Rather, the observed decrease in neural activity during the RF diet is consistent with increased tonic dopamine preferentially engaging inhibitory D2 receptor expressing neurons with a high affinity for dopamine. Stimulation of activity in neurons expressing the lower affinity D1/5 receptor requires phasic dopamine responses ^33^, which are expected to reduce – not increase – dopamine binding at the D2/3 receptor ^34^. Phasic dopamine responses are not expected under our experimental conditions given that the scans were conducted without providing rewards or reward-predicting stimuli. Indeed, visual food stimuli do not result in detectable changes in dopamine in humans ^35,36^. Therefore, our multi-modal neuroimaging results are most likely explained by increases in tonic striatal dopamine resulting from the RF diet.

An increase in tonic dopamine during the RF diet occurred in conjunction with an increased selection of high-fat, high-carbohydrate foods observed during the subsequent exploratory *ad libitum* period. Elevations in tonic dopamine alter the balance with phasic dopamine responses ^33^ and may increase incentive salience ^37,38^, enhance the ‘wanting’ of foods high in both carbohydrate and fat that are particularly rewarding ^39,40^, promote selection of these foods previously experienced to deliver reward ^41^ and reduce the influence of any ensuing negative outcomes on changing behavior ^42^. It is intriguing to speculate about a role for tonic dopamine in influencing the food choices and making it difficult for people to adhere to low fat diets, at least in the short term.

At first glance, our observation that a reduction in BOLD response to food cues during the RF diet occurred alongside a subsequent shift in ad libitum food selections toward high-fat, high-carbohydrate foods appears at odds with the literature on food cue reactivity suggesting a moderate positive association with subsequent weight gain and eating behavior ^43^. However, previous studies employed cross-sectional designs in participants consuming their habitual diets and did not experimentally manipulate the BOLD response to food cues. We speculate that decreased striatal BOLD response during the RF diet may be due to increases in tonic dopamine engaging inhibitory D2 receptor expressing neurons thereby biasing food choice towards rewarding foods.

How could reduction of dietary fat result in increased tonic dopamine in the brain? Dietary fat is detected and signaled to the brain throughout the alimentary canal from tastebud cells in the oral cavity to enteroendocrine and enterocyte cells in the gut ^44^. One of several mechanisms by which dietary fats modulate feeding includes intestinal production of oleoylethanolamide (OEA), a lipid messenger produced from dietary oleic acid which can signal to the brain via the vagus nerve ^45-49^. Despite OEA being produced from dietary fat, chronic consumption of high fat diets in rodents decreases intestinal production of OEA and decreases brain dopamine ^48^. Perhaps because our study participants with established obesity reduced their fat intake by ∼90 grams per day during the RF diet, their intestinal OEA production may have increased thereby resulting in increased brain dopamine. In that case, the effect of increased intestinal OEA production might be expected to enhance satiety during the RF diet ^45-49^ while at the same time the increased tonic dopamine might have steered food choices away from such a diet towards more rewarding foods. In other words, adhering to a low-fat diet might be difficult despite it potentially being more satiating and leading to decreased *ad libitum* energy intake in a setting where “off diet” foods are unavailable ^50^.

Another potential mechanism for increased brain dopamine during the RF diet involves decreased postprandial plasma triglycerides that peak several hours after a meal in proportion to the amount of fat consumed ^51^. Triglycerides have been shown to suppress dopamine synthesis and excitability of D2/3 receptor expressing neurons ^52^ as well as to influence the preference for palatable food and reward seeking in mice ^53^. Compared to the baseline and RC diets, the RF diet would be expected to result in reduced postprandial triglycerides and therefore increased brain dopamine at the times of the neuroimaging scans conducted 2-3 hours postprandially.

Why did the RC diet have no effect on brain D2BP or neural activity in response to food cues as compared to baseline? We found this result surprising particularly because the RC diet significantly decreased daily insulin secretion ^2^ and would be expected to decrease insulin in the brain ^54^, influencing multiple aspects of the dopamine system. For example, dopaminergic neurons express insulin receptors ^55^ and insulin decreases synaptic dopamine by increasing clearance from striatal synapses via enhanced dopamine transporter activity ^56,57^. Consistent with a decrease in synaptic dopamine, intranasal insulin delivery was recently observed to increase D2BP in humans ^21^. Therefore, the lack of effect of the RC diet on brain dopamine remains a mystery. Whereas previous studies have demonstrated that calorie restriction potentiates dopaminergic signaling in both rodents and humans ^58,59^, our results using 30% calorie restricted RC and RF diets suggest that restriction of dietary fat may have a more potent effect on brain dopamine than isocaloric restriction of carbohydrates.

How might changes in brain dopamine in response to different diets relate more generally to body weight regulation? Recent mouse data suggest that the effects of brain dopamine may not be isolated to canonical hedonic pathways of food reward. For example, striatal dopamine can also influence downstream hypothalamic nuclei traditionally attributed to control homeostatic feeding and regulate body weight ^3,60^ ultimately promoting intake of foods that cause obesity and devaluing foods that do not result in obesity ^60^. It is therefore intriguing to speculate that diet composition may contribute to altering the homeostatic body weight “set point” via changes in brain dopamine.

## LIMITATIONS

While our interpretation of increased tonic dopamine is supported by relative pharmacokinetic properties of D1/5 and D2/3 receptors, and literature on D2/3 receptor PET occupancy and fMRI activity, we did not directly measure brain dopamine. Consumption of dietary fat elicits rapid dopaminergic response in reward regions ^61,62^. While we observed an effect of reduced fat diet at the D2 receptor via tonic levels of dopamine, it is possible that the relatively high proportion of dietary fat in the reduced carbohydrate diet may have influenced DA system via mechanisms dependent on D1/5 receptors not examined here. Future studies are needed to delineate the effect of exposure duration (single meal versus multi-day), receptor subtype-specific effects (availability of D1/5 versus D2/3 receptors after exposure), and subsequent effect on *ad libitum* eating behavior.

*Ad libitum* eating behavior subsequent to the five-day period of dietary restriction supports our interpretation of increased incentive salience for rewarding foods after the RF diet. However, our study was not specifically powered to detect differences in this exploratory outcome and analyses were not corrected for multiple comparisons.

Our interpretation on the effect of RC and RF diets on brain dopamine is limited to the early stages of initiating reduced energy diets and does not address long term changes or adaptations in neurochemistry or reward. Future studies should investigate changes in neurochemistry and reward activity in relation to diet composition over longer periods of weight loss.

Finally, the number of participants completing neuroimaging scans is relatively small. The power and measurement reliability supplied by this sample size was greatly enhanced, however, by a within-subject random order crossover study designed to test the effect of specific dietary interventions relative to participants’ own brain at baseline ^63^. Nevertheless, our findings warrant future replication.

## ACKNOWLEDGEMENTS

This work was supported by the Intramural Research Program of the National Institutes of Health, National Institute of Diabetes and Digestive and Kidney Diseases. We thank the nursing and nutrition staff at the NIH MCRU for their invaluable assistance with this study. We are most thankful to the study subjects who volunteered to participate in this demanding protocol.

## METHODS

### Experimental Model and Subject Detail

Twenty-one adults provided informed consent to participate in a randomized crossover trial investigating the effects of selective isocaloric reduction of dietary fat versus carbohydrate on macronutrient metabolism, striatal dopamine type 2 receptor binding potential, and neural activity in response to food stimuli in brain reward regions (ClinicalTrials.gov NCT00846040). Study details regarding the primary metabolic outcomes were reported elsewhere (Hall et al., 2015). In brief, right-handed non-smokers between 18-45 years of age with a reported BMI greater than 30 kg/m^2^ (body weight < 350 pounds) were recruited from the Washington DC metro area. All were free from diabetes, recent weight change (> ± 5 kg in the past 6 months), physical mobility impairments, past or present history of drug abuse, neurological or psychiatric disorders (including eating disorders such as binge eating) as assessed by an abbreviated Structured Clinical Interview for the Diagnostic and Statistical Manual of Mental Disorders. Furthermore, participants were free from evidence of diseases or medications interfering with study outcomes, allergies to food or local anesthetics, evidence of regular excessive use of caffeinated drinks and alcohol or strict dietary concerns (vegetarian or kosher diet). Premenopausal women were studied in the follicular phase for each inpatient visit and were excluded if they were pregnant or breastfeeding. All study procedures were approved by the Institutional Review Board of the National Institute of Diabetes & Digestive & Kidney Diseases and participants were compensated for their participation.

### Method Details

Volunteers were admitted to the NIH Clinical Center for a 14-day period to receive the eucaloric baseline diet for 5 days followed by either the RC or the RF diet for the next 6 days followed by 3 days of *ad libitum* feeding from a computerized vending machine, as detailed below (Figure 1). Participants were readmitted after a 2-to 4-week washout period to repeat the 5-day eucaloric baseline diet followed by 6 days of the alternate reduced calorie diet and 3 days of *ad libitum* feeding. Every day, participants completed 60 min of treadmill walking at a fixed self-selected pace and incline determined during screening to mimic free-living levels of physical activity.

The CONSORT diagram reiterates enrollment details provided in ^2^ (Supplementary Figure 1). Two participants withdrew during the first baseline diet and did not complete any neuroimaging. Of the 19 participants who completed the initial baseline diet, 10 were randomized to next receive the RC diet and 9 were randomized to next receive the RF diet. Among 10 participants receiving the RC diet on their first admission, 1 participant completed PET but not fMRI procedures during the RC diet and 2 withdrew before receiving the RF diet on the second planned admission. Among the 9 participants receiving the RF diet on their first admission, 2 completed fMRI but not PET on their first admission (participant-declined PET scans), and 1 participant did not have available fMRI data during their second admission on the RC diet. Full neuroimaging data (PET and fMRI) across all 3 diet conditions are available for n=13 participants and the results are provided in Supplementary Materials. Complete PET data are available in n=15 participants and complete fMRI data are available from n=15 participants and the results are provided in Supplementary Materials.

### Anthropometrics

Height was measured in centimeters using a wall stadiometer (Seca 242, Hanover, MD, USA) and weight was measured in kilograms using a digital scale (Scale-Tronix 5702, Carol Steam, IL, USA). All measurements were obtained after an overnight fast while participants were wearing only hospital scrubs.

### Diets

All subjects were confined to the metabolic ward throughout the study without access to outside food. Meals were consumed under observation and any uneaten food was returned to the kitchen and re-weighed. Subsequent meals were adjusted to account for uneaten food as needed. Diets were designed using ProNutra software (version 3.4, Viocare, Inc.).

#### Baseline eucaloric diet

The daily caloric content during the initial out-patient segment and the weight-maintenance phase was based on the resting energy expenditure measured at screening with an activity factor of 1.5. Beginning 2 days before each admission, participants were provided with a weight-maintenance diet using a standard diet composition of 50% carbohydrate, 35% fat, and 15% protein, which continued for the next 5 days. All participants were provided with the standard diet during the first inpatient admission for at least one day prior to measuring baseline fMRI and D2BP. Energy and macronutrient intake during the baseline eucaloric diet are presented in Table 1.

#### Reduced energy diets

During the restricted diet phase (inpatient days 6 to 11), 30% of baseline calories were removed by selective reduction of either carbohydrate (RC diet) or fat (RF diet) while keeping the other two macronutrients unchanged from eucaloric baseline diet. Energy and macronutrient intake during the reduced energy diets are presented in Table 1.

#### *Ad libitum* vending machine diet

For the last 3 days of each inpatient stay, participants were given *ad libitum* access to a computerized vending machine (Starfoods, Necta, Valbrembo, Italy). The MSSP procedure was used to select items for stocking the vending machine ^64^. This paradigm was selected for use in this study as it was developed to show preference between foods of differing fat and carbohydrate content. It is composed of six categories of food, including high complex carbohydrate/high fat (HCHF), high simple sugar /high fat (HSHF), low carbohydrate/high protein/high fat (HPHF), high complex carbohydrate/low fat (HCLF), high simple sugar/low fat (HSLF), and low carbohydrate/high protein/high fat/ (HPHF). A list of 76 foods that fit into these categories was provided to participants in a Food Questionnaire. This questionnaire contained Likert-type scales with questions in which the participant rated how much they liked each of the food items that could potentially be provided in the vending machine. The questionnaire also asked how often each of those food items were consumed normally by the participant (i.e., daily, weekly, or monthly). Of these foods, a total of 40 items that fit into the previously mentioned categories were chosen for inclusion in the vending machine if preference was rated from 4 to 9 on the 10-point Likert scale.

Vending machines were stocked with traditional breakfast, lunch, dinner and snack items. Beverages and condiments were also included in the vending machine and consumption of these items was also recorded. Sugar-sweetened beverages (SSB) included fruit juices, lemonade, chocolate milk and regular sodas. Each participant had access to one vending machine that only they could access. Once foods were selected, participants were instructed to eat in the dining area and no food was allowed in the participant’s room. All uneaten food and wrappers were returned to the Metabolic Kitchen to be weighed. The vending machines were re-stocked daily at 8 am with items that had been removed in the previous 24 hours. All foods were weighed to the nearest tenth of a gram on a digital scale (Mettler Toledo MS Series, Columbus, OH, USA) prior to placing them in the computerized vending machine and the remainder of any uneaten foods were weighed after consumption. Energy and macronutrient composition of the foods consumed from the vending machine were calculated using a computerized nutrition database (ProNutra, Viocare Inc., Princeton, NJ, USA).

Vending machine foods were retrospectively categorized as either ultra-processed or non-ultra-processed based on NOVA categories ^65^ and additionally categorized as hyperpalatable or non-hyperpalatable based on definitions presented by ^66^.

Statistical analyses of caloric intake from Vending Machines were performed using IBM SPSS Statistics (28.0.1.1). Repeated-measure mixed model analyses were used to assess differences in intake of energy, macronutrients, percent of calories from MSSP and sugar sweetened beverages among 17 participants completing both 3-day *ad libitum* periods.

### Magnetic Resonance Imaging

On the afternoon following the morning PET scanning, high resolution anatomical brain MRI was acquired with a HDx General Electric 3 Tesla scanner (TE = 2.7ms, TR 7.24 ms, flip angle 12°, voxel size 0.937*0.937*1.2mm) for each subject.

Under each diet condition, all subjects were scanned at 18:00, 4.5 h after a standardized, diet-appropriate meal. Functional and structural imaging was performed on a 3T General Electric scanner and a GE 8-channel receive-only head coil. High-resolution anatomical images were collected prior to functional scanning runs (TE = 2.7 ms, TR: 7.24 ms, flip angle: 12 degrees, voxel size: 0.937×0.937×1.2 mm). For the functional scans, 206 magnetic resonance (MR) volumes were acquired. Each echoplanar image (EPI) consisted of 44 2.8-mm slices (echo time [TE] = 27 ms, repetition time [TR] = 2500 ms, flip angle = 90 degrees, voxel size = 3.4375×3.4375×2.8 mm). All structural and functional images were collected with a Sensitivity Encoding (SENSE) factor of 2 used to reduce image collection time (for structural images) or minimize image distortions (in functional images) while reducing gradient coil heating over the course of the scan session.

The fMRI task is described in detail elsewhere (Simmons et al., 2014). In brief, 144 visual food cues ranging from highly processed, energy dense foods to raw fruits and vegetables were displayed to participants using E-prime software (www.pstnet.com). Images projected to the scanner-room screen were viewed via head coil-mounted mirror. Each image was presented for 5 seconds during which time participants indicated their response to a question (“If given the opportunity right now, how pleasant would it be to eat this food?”) using an MR compatible scroll wheel to select values along a number line positioned next to the image. A fixation cross was presented for varying durations between stimuli (mean ISI = 3.7 seconds; duration 2.5–7.5 seconds). The pleasantness rating scale ranged from 1 (“neutral”) to 7, with 1 depicted as “neutral” and 7 as “extremely pleasant” and included an “unpleasant” option represented by the letter “X” located below the number line. For images that participants viewed as “unpleasant”, they were instructed to select the “X” if they believed the depicted food would be at all unpleasant to eat. Food images rated as “unpleasant” were excluded from the MRI and behavioral analyses.

Analyses of functional neuroimaging were performed in AFNI (AFNI_20.2.00 ‘Aulus Vitellius’). Each individual’s anatomical MRI was transformed into the Talairach space, and the transformation matrix was applied to the functional data during pre-processing. All functional volumes were aligned to a common base EPI represented by the third volume of the first functional run. The first three volumes of each EPI run were trimmed to allow the fMRI signal to reach steady state. A slice-time correction was applied to all functional volumes, which were also smoothed with a 6-mm full-width half-max Gaussian kernel. Additionally, the signal value for each EPI volume was normalized to the percent signal change from the voxel’s mean signal across the time course.

Individual subject data were checked for quality assurance, and outlying time points resulting from head motion were censored from the analyses. At the individual level, multiple regression was used to analyze the data, with regressors of non-interest included in the model to account for each run’s signal mean, linear, quadratic, and cubic signal trends, as well as six motion parameters (three translations and three rotations) saved from the image registration step during pre-processing. The food pleasantness task regressor was constructed by convolving a box-car function with a width of 5 s beginning at the onset of the food image with a gamma-variate function to adjust the predictor variable for the delay and shape of the BOLD response. Given similarities in pleasantness ratings across diet conditions (Supplementary Figure 2), task pleasantness ratings were not included as parametric modulators of the hemodynamic response.

### Positron Emission Tomography

PET scanning was performed using a High Resolution Research Tomograph (HRRT; Siemens Healthcare, Malvern, PA) a dedicated brain PET scanner with a resolution of 2.5-3.0 mm and a 25 cm axial field of view. Transmission scanning was performed with a 137Cs rotating pin source to correct for attenuation. Two hours after a standard breakfast, a bolus of approximately 5 mCi of [18F]fallypride was infused intravenously using a Harvard® pump. The specific activity was approximately 2000 mCi/μmol at time of injection and the radiochemical purity of the radiotracer was > 99%. PET emission data were collected starting at radiotracer injection over 3.5 hours, in three blocks separated by two 10-minute breaks. Thirty-three volumes were acquired at times 0, 0.25, 0.5, 0.75, 1, 1.25, 1.5, 1.75, 2, 2.5, 3, 3.5, 4, 4.5, 5, 6, 7, 8, 9, 10, 12.5, 15, 20, 25, 30, 40, 50, 60, 90, 110, 130, 170, 200 min. During each scan block, the room was quiet and dimly lit and each subject was instructed to keep their head as still as possible, relax, and try to avoid falling asleep. The image reconstruction process corrected for head motion which was tracked throughout each scan. Each scan consisted of 207 slices (slice separation = 1.22 mm). The fields of view were 31.2 cm and 25.2 cm for transverse and axial slices, respectively.

The PET images were aligned within each scan block with 6-parameter rigid registration using 7th order polynomial interpolation and each block was aligned to the volume taken at 20 min of the first block. The final alignments were visually checked, with translations varying by <5 mm and the rotations by <5 degrees.

### Quantification and Statistical Analysis

fMRI images were included in AFNI’s 3dttest++ to identify clusters of significant effects of the diet condition (RF>Baseline for n=17; RC>Baseline for n=17; RF>RC for n=15). Analyses using participants with complete neuroimaging data (fMRI and PET) across 3 diet conditions (n=13) were analyzed via AFNI 3dANOVA (Supplementary Materials). Since diet condition did not have a significant impact on food pleasantness ratings, analysis of brain activity to food pictures was not modulated by pleasantness ratings to maximize study power. Small volume corrections were implemented within the ROI defined by the orbitofrontal cortex, striatal-pallidal reward regions as previously described ^20^.with a voxel-wise p<0.001 and cluster size threshold (k_e_>5) to achieve bi-sided correction for multiple comparisons at p<0.05 via AFNI 3dclustsim

Individual participants’ anatomical MRI images (see above) were co-registered to the aligned PET images by minimizing a mutual information cost function for each individual participant. For the analyses described in the main text, each individual’s anatomical MRI was linearly transformed into the Talairach space, and the transformation matrix was applied to the PET images which were then smoothed with a 5-mm full-width, half-max Gaussian kernel. Data were exported to MATLAB where time-activity curves for [18F] fallypride concentration in each voxel were fit to a kinetic model (with the cerebellum used as the reference tissue) to determine D2BP ^67^. In an alternative pipeline presented in the Supplementary Materials, PET images were first smoothed and D2BP was calculated in native space followed by non-linear warping to Talairach space.

Participants’ D2BP maps were included in AFNI’s 3dttest++ identify clusters with significant effects of diet (RF>Baseline for n=15; RC>Baseline for n=17; RF>RC for n=15). Analyses using participants with complete neuroimaging data across 3 diet conditions (n=13) were analyzed via AFNI 3dANOVA (Supplementary Figure 3E). Since high D2BP occurs mainly in striatum, small volume corrections were implemented within each hemisphere where D2BP >1.5. A bi-sided voxel-wise threshold of p<0.1 was used, and cluster size threshold to achieve correction for multiple comparisons at p<0.05. Using a full mixed effects model (AFNI 3dANOVA3), clusters survive correction for multiple comparisons using 3dClustSim at alpha of 0.05 a threshold of 33 voxels.

To test the robustness of our results with respect to alterations in processing and analysis pipeline, we also analyzed the data Individual participants’ anatomical MRI images were co-registered to the aligned PET images by minimizing a mutual information cost function for each individual participant. The aligned PET images were smoothed with a 5-mm full-width, half-max Gaussian kernel. Data were exported to MATLAB where time-activity curves for [18F] fallypride concentration in each voxel were fit to a kinetic model with the cerebellum used as the reference tissue to determine D2BP ^67^. The values of D2BP were then imported back into individual native spaces to construct D2BP maps. Each individual’s anatomical MRI image was mapped into the Talairach space with the AFNI program auto_warp.py and produced a non-linear transformation function, which was then applied to transform each individual D2BP maps into the Talairach space (Supplementary Figures 5B and 6B).

