## Supplementary Information for "Restriction of dietary fat, but not carbohydrate, affects brain reward regions in adults with obesity"

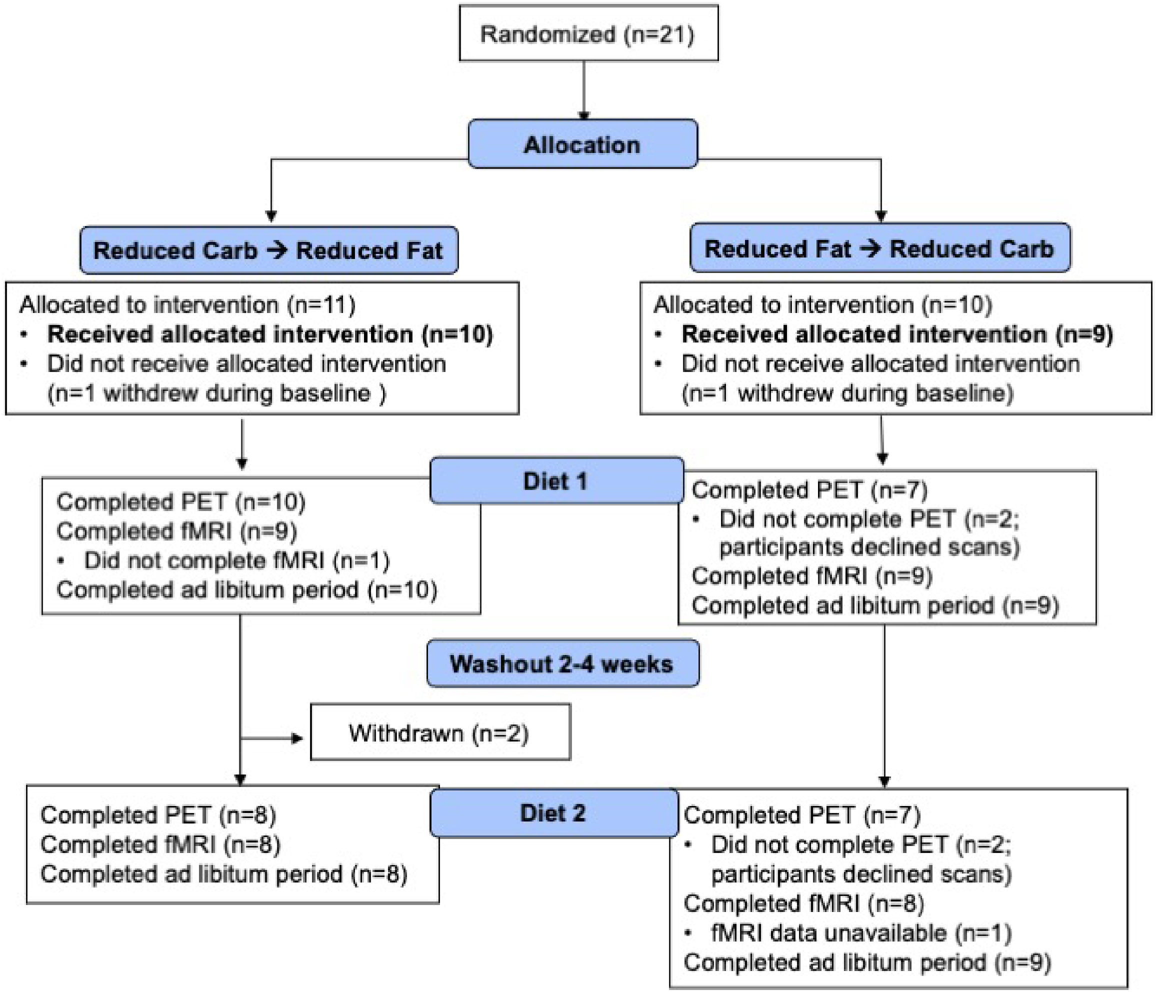


**Supplementary Figure 1. CONSORT diagram of participant randomization in crossover trial.** Full neuroimaging data (PET and fMRI) across all 3 diet conditions are available for n=13 participants and the results are provided in Supplementary Materials. Complete PET data across 3 diet conditions are available in n=15 participants and complete fMRI data across 3 diet conditions are available from n=15 participants.

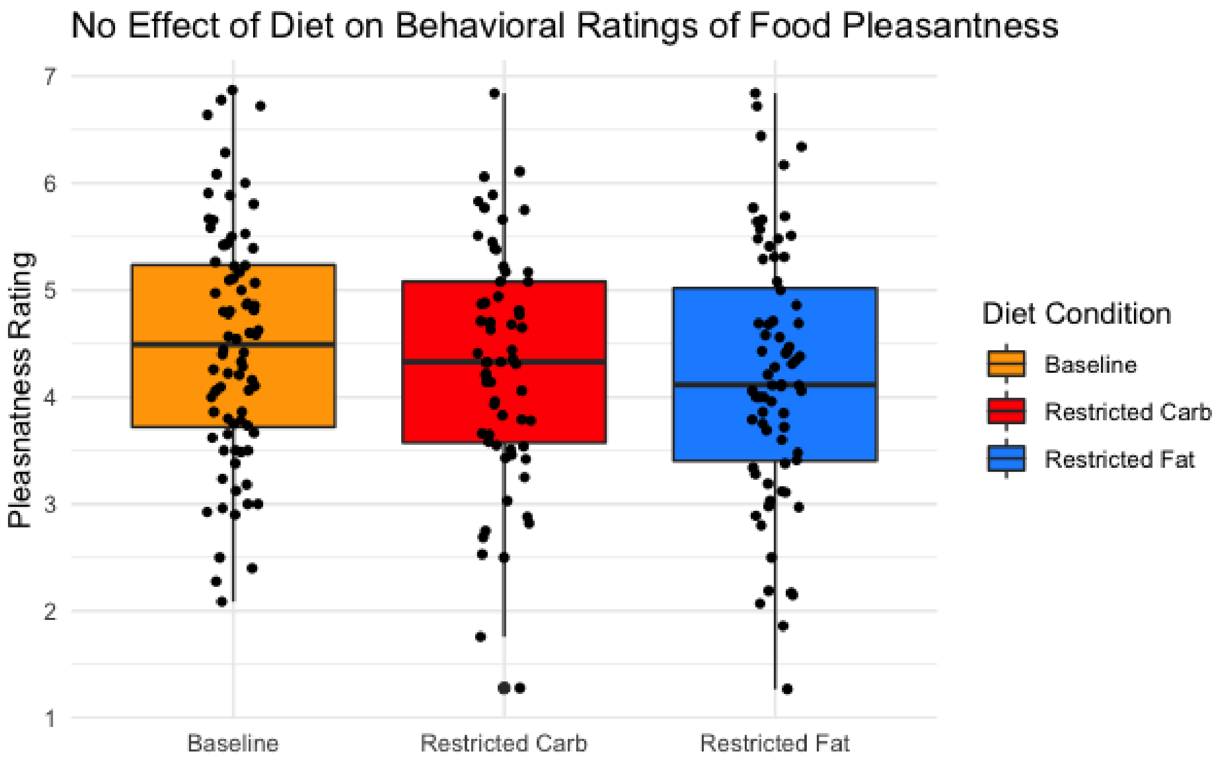


**Supplementary Figure 2. Diet condition did not significantly impact in-scanner pleasantness ratings of visual food cues.** N=15; Means: 4.39 ± 3.7 at Baseline, 4.35 ± 0.58 at RC, 4.30 ± 0.58 at RF; Repeated measures ANOVA Wilks' Lambda 0.29, *F* 0.498, *p*=0.619.

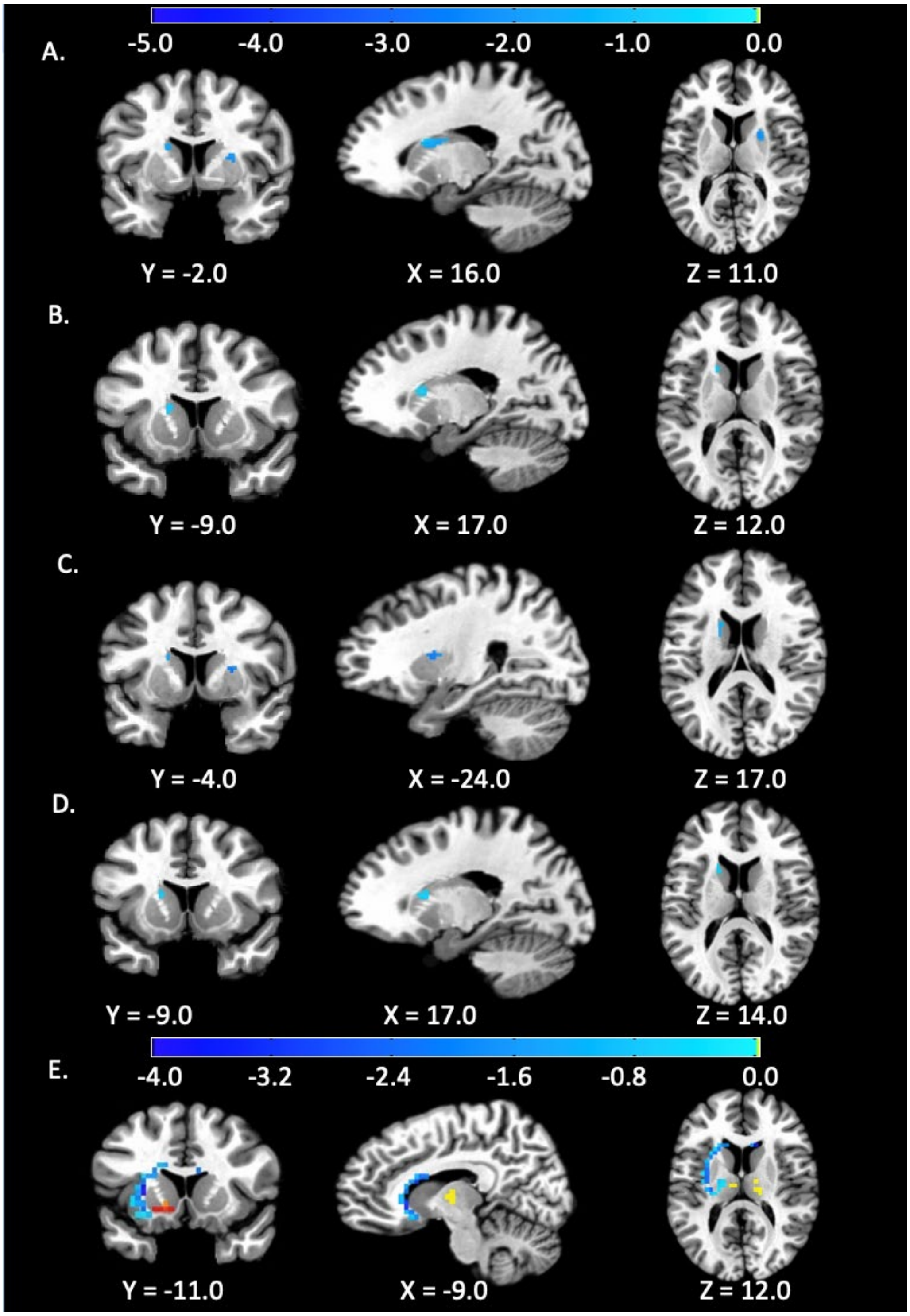


**Supplementary Figure 3. Reduced-fat diet decreased both striatal activity in response to visual food cues and dopamine D2BP.** fMRI analyses including 15 participants with complete fMRI data, contrasting (A) RF versus Baseline diets and (B) RF versus RC diets. fMRI analyses including 13 participants with both complete fMRI and PET data, contrasting (C) RF versus Baseline diets, and (D) RF versus RC diets. (E) Analysis of D2BP in 13 participants with complete fMRI and PET data, contrasting RF versus baseline diets. See Supplementary Table 1 for corresponding cluster details.

**Supplementary Table 1. Locations of clusters displaying changes in BOLD responses within *a priori* reward region mask or D2BP to reduced fat or reduced carbohydrate diet within individuals completing all diets and all fMRI (n=15), all diets and all PET scans (n=15), and all diets, fMRI and PET (n=13).**

| | *Location of peak* | | | *Voxels* | *Size ($mm^3$)* | *Z-score (fMRI) t-stat (PET)* | *alpha* |
|---|---|---|---|---|---|---|---|
| | *x* | *y* | *z* | | | | |
| **<u>fMRI</u>** | | | | | | | |
| **Reduced Fat vs. Reduced Carbohydrate v. Baseline** | | | | | | | |
| **(**ANOVA, n=15 with complete fMRI; $k_e$=5, $p_{uncorrected}$ = 0.001) | | | | | | | |
| *RF diet vs. Baseline* | | | | | | | |
| Left caudate | 15.0 | 1.0 | 18.0 | 38 | 304 | -3.43 | <0.01 |
| Right putamen | -23.0 | -5.0 | 10.0 | 21 | 168 | -3.31 | <0.02 |
| Right mid. orbital gyrus | -25.0 | -45.0 | -10.0 | 6 | 48 | -3.38 | <0.09 |
| Right ventromedial putamen | -23.0 | -5.0 | 0.0 | 6 | 48 | -3.51 | <0.09 |
| *RF diet vs. RC diet* | | | | | | | |
| Left caudate | 17.0 | -9.0 | 12.0 | 16 | 128 | -3.33 | <0.03 |
| *RC diet vs. Baseline* | | | | | | | |
| No clusters | -- | -- | -- | -- | -- | -- | -- |
| **Reduced Fat vs. Reduced Carbohydrate v. Baseline** | | | | | | | |
| (ANOVA, n=13 with complete PET & fMRI; $k_e$=5, $p_{uncorrected}$ = 0.001) | | | | | | | |
| *RF diet vs. Baseline* | | | | | | | |
| Right putamen | -23.0 | -3.0 | 10.0 | 17 | 136 | -3.30 | <0.02 |
| Left caudate | 17.0 | -7.0 | 16.0 | 18 | 144 | -3.34 | <0.02 |
| *RF diet vs. RC diet* | | | | | | | |
| Left caudate | 17.0 | -9.0 | 14.0 | 11 | 88 | -3.45 | <0.04 |
| *RC diet vs. Baseline* | | | | | | | |
| No clusters | -- | -- | -- | -- | -- | -- | -- |
| **<u>PET D2BP</u>** | | | | | | | |
| **Reduced Fat vs. Reduced Carbohydrate v. Baseline** | | | | | | | |
| (ANOVA, n=15 with complete PET**;** $k_e$=20, $p_{uncorrected}$ = 0.1) | | | | | | | |
| *RF diet vs. RC diet* | | | | | | | |
| No clusters | -- | -- | -- | -- | -- | -- | -- |
| *RC diet vs. Baseline* | | | | | | | |
| No clusters | -- | -- | -- | -- | -- | -- | -- |
| **Reduced Fat vs. Reduced Carbohydrate v. Baseline** | | | | | | | |
| (ANOVA, n=13 with complete PET & fMRI **;** $k_e$=20, $p_{uncorrected}$ = 0.1)**)** | | | | | | | |
| *RF diet vs. Baseline* | | | | | | | |
| Left putamen | 26.2 | -9.5 | 6.5 | 172 | 7375 | -2.33 | <0.01 |
| Right caudate | -12.2 | -20.0 | 13.5 | 50 | 2144 | -1.93 | <0.01 |
| Right thalamus | -1.8 | 18.5 | 3.0 | 34 | 1457 | 2.87 | <0.05 |
| Left putamen | 19.2 | -6.0 | -4.0 | 26 | 1115 | 3.78 | <0.14 |
| *RF diet vs. RC diet* | | | | | | | |
| No clusters | -- | -- | -- | -- | -- | -- | -- |
| *RC diet vs. Baseline* | | | | | | | |
| No clusters | -- | -- | -- | -- | -- | -- | -- |

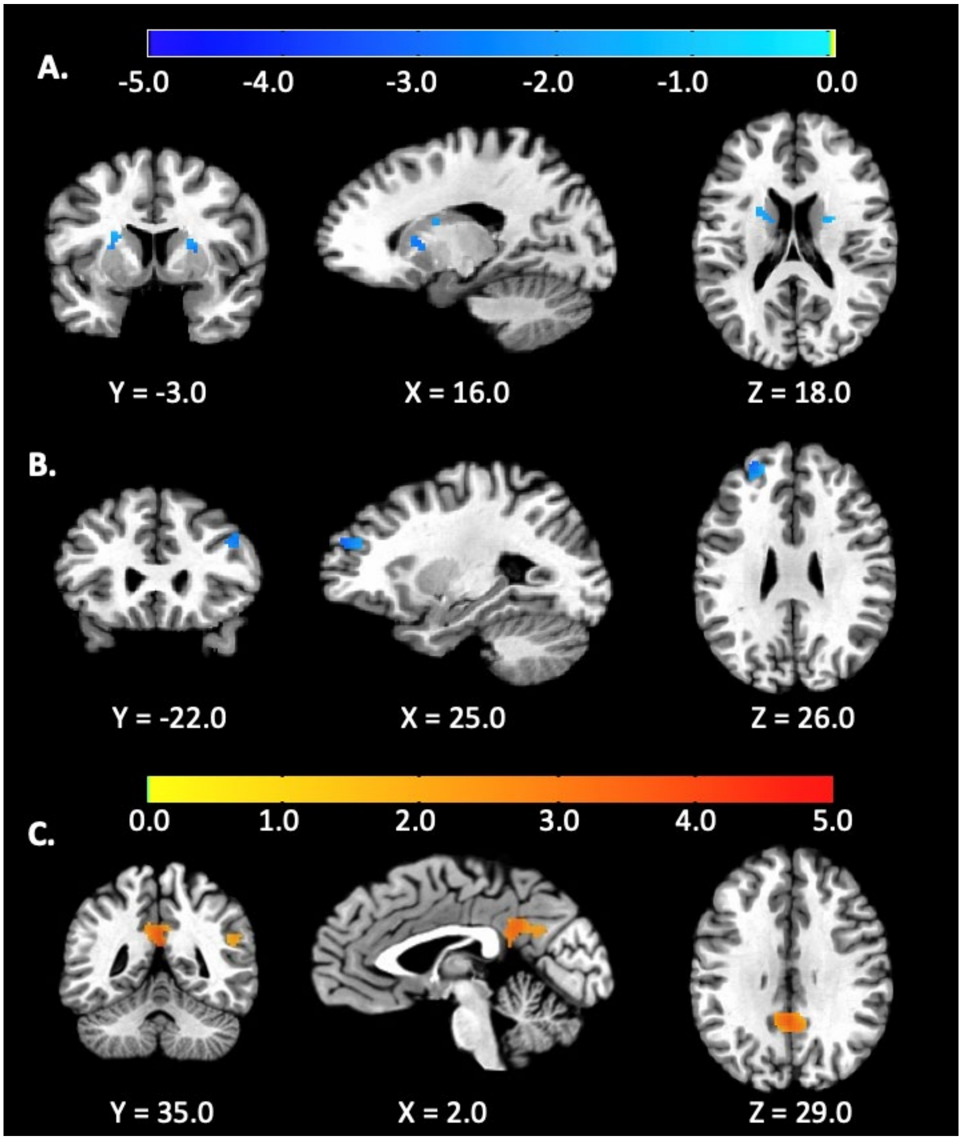


**Supplementary Figure 4. Whole-brain fMRI analyses.** fMRI analyses contrasting (A) RF versus Baseline (n=17), (B) RF versus RC (n=15), and (C) RC versus Baseline (n=17).

**Supplementary Table 2. Locations of clusters displaying changes in whole-brain BOLD responses to reduced fat or reduced carbohydrate diets.**

| | *Location of peak* | | | *Voxels* | *Size ($mm^3$)* | *Z-score* | *alpha* |
|---|---|---|---|---|---|---|---|
| | *x* | *y* | *z* | | | | |
| **Reduced Fat v. Reduced Carbohydrate v. Baseline** | | | | | | | |
| *RF diet vs. Baseline* (*paired t-test, n=17; $k_e$=40, $p_{uncorrected}$ = 0.001)* | | | | | | | |
| Left putamen | 19.0 | -9.0 | 6.0 | 77 | 616 | -3.48 | <0.06 |
| Right putamen | -23.0 | 3.0 | 10.0 | 50 | 400 | -3.41 | >0.10 |
| *RC diet vs. Baseline* (*paired t-test, n=17; $k_e$=40, $p_{uncorrected}$ = 0.001)* | | | | | | | |
| Right posterior cingulate/precuneus | -1.0 | 45.0 | 20.0 | 234 | 1872 | 3.33 | <0.01 |
| Right middle temporal gyrus | -49.0 | -1.0 | -22.0 | 80 | 640 | 3.80 | <0.07 |
| Left middle temporal gyrus | 61.0 | 31.0 | 0.0 | 64 | 512 | 3.58 | <0.10 |
| Right supramarginal gyrus | -49.0 | 45.0 | 26.0 | 58 | 464 | 3.45 | >0.10 |
| Right angular gyrus | -39.0 | 57.0 | 26.0 | 54 | 432 | 3.79 | >0.10 |
| Right medial temporal pole | -37.0 | -3.0 | -24.0 | 46 | 368 | 3.56 | >0.10 |
| *RF diet vs. RC diet* (*paired t-test, n=15; $k_e$=40, $p_{uncorrected}$ = 0.001)* | | | | | | | |
| Left superior frontal gyrus | 25.0 | -51.0 | 26.0 | 72 | 576 | -3.49 | <0.07 |
| Right inferior frontal gyrus | -45.0 | -15.0 | 32.0 | 42 | 336 | -3.34 | >0.10 |

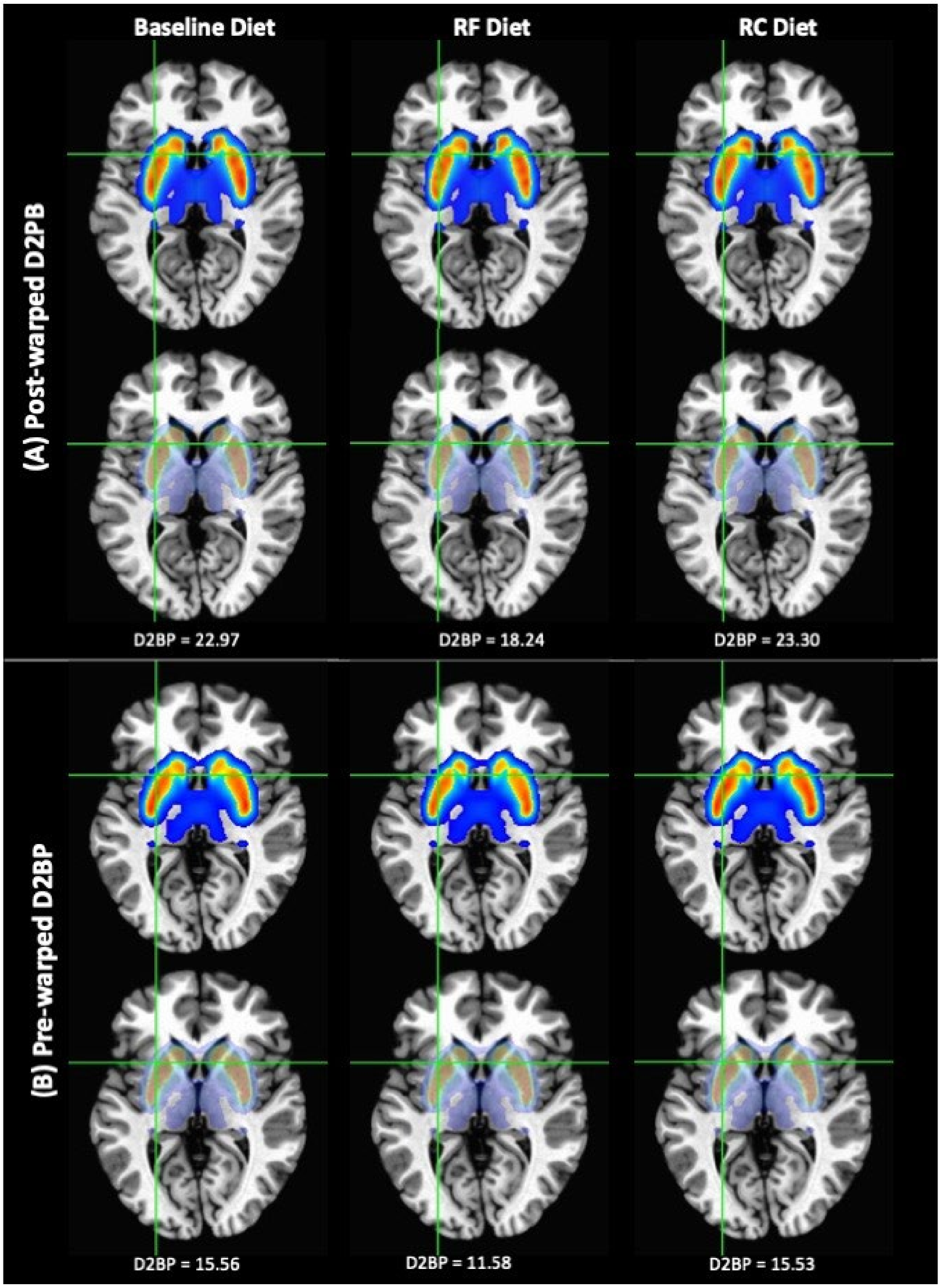


**Supplementary Figure 5. Mean D2BP maps by diet condition in each column**. (A) Data processed according to the pipeline reported in the main text where D2BP was computed after linear warping individual subject data to Talairach coordinates. Top row indicates the mean D2BP>1.5 overlaid on the template brain and the bottom row depicts the results with increased transparency of the D2BP map to visualize anatomical overlap. Peak D2BP was located within striatal nuclei, but regions with nonzero D2BP were found outside these regions, including in white matter. Crosshairs indicate coordinates within the cluster defining maximum significant differences between RF and Baseline diets indicating that this occurred at regions close to gray/white border (26.2, -9.5, 6.5). D2BP at this location for each diet condition is indicated below each column. (B) Data processed according to a pipeline where D2BP was first computed in native space and then nonlinear warped to Talairach coordinates and produces similar results for the location of the cluster defining maximum significant differences between RF and Baseline diets (26.2, -9.5, 3).

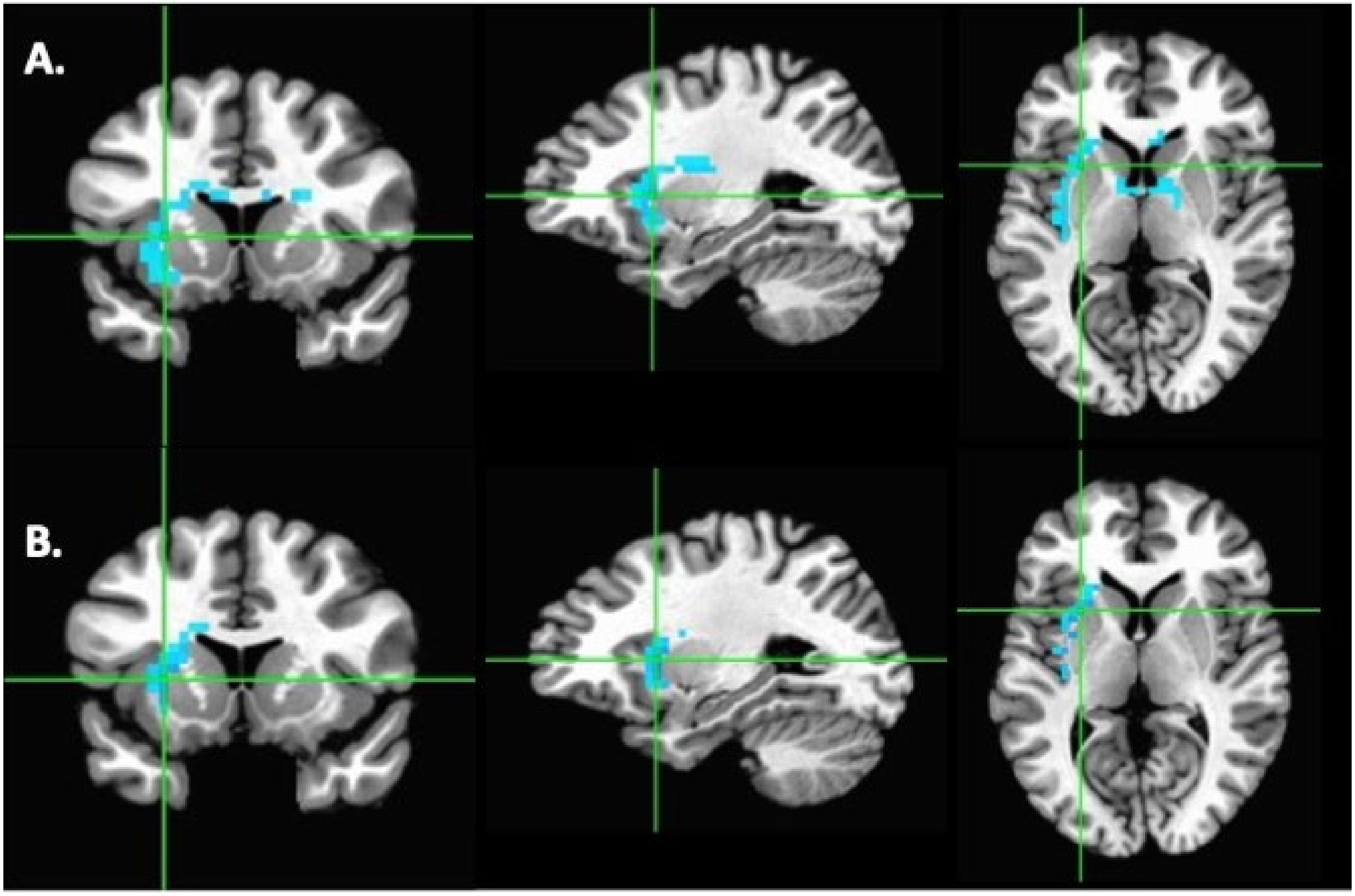


**Supplementary Figure 6.** D2BP contrast between RF and Baseline diets (n=15) using D2BP calculated using (A) a processing pipeline where D2BP was computed after linear warping individual subject data to Talairach coordinates (peak: 26.2, -9.5, 6.5) and (B) a pipeline where D2BP was computed in native space and then nonlinear warped to Talairach coordinates (peak: 26.2, -9.5, 3), producing a similar cluster.
